# A Small-Bodied Troodontid (Dinosauria, Theropoda) from the Upper Cretaceous Wulansuhai Formation of Inner Mongolia, China

**DOI:** 10.1101/2020.02.05.936526

**Authors:** Shuo Wang, Qingwei Tan, Qiyue Zhang, Josef Steigler, Huitao Zhang, Lin Tan

## Abstract

A new small-bodied troodontid (LH PV39) recovered from the Upper Cretaceous Wulansuhai Formation, Suhongtu, Inner Mongolia, China, is described. The new specimen preserves six postaxial cervical vertebrae, five completely fused sacral and four posterior caudal vertebrae in addition to two manual unguals. The completely fused neurocentral junctions indicate that a skeletally mature individual of the same species of LH PV39 would be smaller than *Philovenator* and comparable in body size to a skeletal mature individual of *Almas*. The extremely dorsoventrally compressed sacral centra and neural canal, and the middle three sacral centra that are shorter and wider than the first and the last one distinguishing LH PV39 from other known troodontids. A series of phylogenetic analyses were conducted using modified published matrices. By coding LH PV39 in different strategies, the troodontid affinity of LH PV39 is confirmed and it was recovered as the sister taxon of either *Mei* and *Sinovenator* (LH PV39 scored as a separate OTU) or *Linhevenator* (incorporating LH PV39 into *Philovenator*) in the best resolved coelurosaurian interrelationships. The referral of LH PV39 to *Philovenator* does not seriously alter the phylogenetic position of *Philovenator* nor the interrelationships of troodontids. This new finding confirms that the small and large sized troodontids are coexisted in the Gobi Desert of the Mongolia Plateau until the end of Cretaceous.

---

Troodontids, known primarily from Upper Jurassic through end Cretaceous deposits of North American and Asia, are a group of bird-like small theropods characterized by enlarged braincases, lightly built snouts, and elongate legs, among other characters (Holtz 2012; Makovicky & Norell 2004). Following their initial discovery, the rarity of troodontid discoveries and their fragmentary remains resulted in more than a century of confusion regarding their taxonomy and relationships to other theropod dinosaurs including birds (Gilmore 1924; Horner & Weishampel 1988; Horner & Weishampel 1996; Lambe 1902; Leidy 1856; Norell *et al*. 1994). This confusion has been largely clarified by discoveries of relatively complete troodontids from the Upper Jurassic-Lower Cretaceous deposits of eastern China (Gao *et al*. 2012; Shen *et al*. 2017a; Shen *et al*. 2017b; Xu *et al*. 2017; Xu & Norell 2004; Xu *et al*. 2002; Xu & Wang 2004a), and the Upper Cretaceous deposits from Mongolian Plateau (Barsbold *et al*. 1987; Bever & Norell 2009; Currie & Dong 2001; Kurzanov & Osmólska 1991; Makovicky *et al*. 2003; Norell *et al*. 2009; Norell *et al*. 2000; Osborn 1924; Pei *et al*. 2017b; Tsuihiji 2014; Xu *et al*. 2011a; Xu *et al*. 2012). Troodontids yielded from the Middle-Late Jurassic Yanliao Biota and Early Cretaceous Jehol Biota have significantly broadened our understanding of the early evolution and diversification of this clade (Gao *et al*. 2012; Shen *et al*. 2017a; Shen *et al*. 2017b; Xu *et al*. 2017; Xu & Norell 2004; Xu *et al*. 2002; Xu & Wang 2004a), and the presence of feathers in some taxa provide convincing evidence of their close relationship with birds (Ji 2005; Xu *et al*. 2017). On the other hand, specimens recovered from Upper Cretaceous sediments have enriched the taxonomic diversity and morphological disparity of Late Cretaceous theropods (Barsbold *et al*. 1987; Bever & Norell 2009; Currie & Dong 2001; Kurzanov & Osmólska 1991; Makovicky *et al*. 2003; Norell *et al*. 2009; Norell *et al*. 2000; Osborn 1924; Pei *et al*. 2017b; Tsuihiji 2014; Xu *et al*. 2011a; Xu *et al*. 2012), making up for the relatively rare materials of troodontids from the contemporaneous sediments in other continents (Evans *et al*. 2017).

Late Cretaceous troodontid taxa are usually larger-bodied than those collected from older sediments (Xu *et al*. 2012), suggesting that one or more instances of gigantism characterize the evolution of troodontids, as in deinonychosaurians generally (Turner *et al*. 2007; Xu *et al*. 2012). However, exceptionally small body size makes *Almas ukhaa* (Pei *et al*. 2017b) from the Late Cretaceous Djadokhta Formation of Ukhaa Tolgod, Mongolia, and *Philovenator curriei* (Xu *et al*. 2012) from the Wulansuhai Formation of Inner Mongolia at Bayan Mandahu stand out from other Late Cretaceous troodontids (Bever & Norell 2009; Currie & Dong 2001; Currie & Peng 1993; Kurzanov & Osmólska 1991; Makovicky *et al*. 2003; Norell *et al*. 2009; Norell *et al*. 2000; Osborn 1924; Tsuihiji 2014; Xu *et al*. 2011a; Zanno *et al*. 2011). Known from a single left hindlimb, *Philovenator curriei* was originally identified as a juvenile *Saurornithoides mongoliensis* on the basis of its size (Currie & Peng 1993), but it was recently reidentified as a new small bodied taxon related to *Linhevenator* based on a histological analysis (Xu *et al*. 2012). Here, we report a new troodontid specimen recovered from the Upper Cretaceous Wulansuhai Formation (Cenomanian), Suhongtu area, Inner Mongolia, China, which resembles the sympatric *Philovenator curriei* in its small body size. This new specimen was recovered during the 2001 installment of the Chinese-American joint expedition, and has not yet been studied in detail. Although there is no available limb bones for histological examination, the completely closed neurocentral sutures across the axial column indicate the mature status of the specimen. This new finding confirms that small bodied troodontids do exist in the Upper Cretaceous Wulansuhai Formation, and emphasizes the size disparity among Later Cretaceous troodontids would have been more significant than previously expected.

## Methods for Mi-CT scan

The samples were placed approximately 20 cm from the source and 40 cm from the detector. The resolutions of the Mi-CT images are 160μm, and a total of 720 transmission images were required for each sample, and were reconstructed in a 2048*2048 matrix of 1563 slices using two-dimensional reconstruction software developed by the Institute of High Energy Physics, Chinese Academy of Sciences. The 3D reconstruction was performed using Mimics (Version 15.0).

### Institutional Abbreviations

**DLXH,** Dalian Xinghai Museum, Dalian, China; **DNHM,** Dalian Natural History Museum, Dalian, China; **MPC-D** (formerly known as **IGM**, Institute of Geology, Mongolia), Institute of Paleontology, Mongolian Academy of Sciences, Ulaanbaatar, Mongolia; **IVPP,** Institute of Vertebrate Paleontology and Paleoanthropology, Chinese Academy of Sciences; **LH,** Long Hao Institute of Geology and Paleontology, Hohhot, China; **MOR,** Museum of the Rockies, Bozeman, USA.

### Systematic Paleontology

Theropoda Marsh, 1881

Coelurosauria Huene, 1920

Maniraptora Gauthier, 1986

Troodontidae Gilmore, 1924 Genus and species indet.

### Materials

LH PV39, partial postcranial skeleton, including six cervical, five sacral, and four caudal vertebrae that are probably from the middle part of the tail, and two manual unguals.

### Locality and horizon

Upper Cretaceous (Campanian) Wulansuhai Formation, Suhongtu, Alashanzuo Banner, Inner Mongolia, China.

### Description

the preserved axial column is represented by 6 postaxial cervical vertebrae (including a partial prezygapophysis in articulation with the parapophysis), 5 completely fused sacral and 4 posterior caudal vertebrae. All elements have been prepared out of the matrix in three dimensions (Figs. 1-3). Other material of LH PV39 includes two manual unguals, one of which is nearly complete and the other is fragmentary (Figs. 1 & 3). Note throughout the paper, the nomenclatures of vertebral structures are based on Wilson *et al*., 2009; 2011.

**Figure 1.**
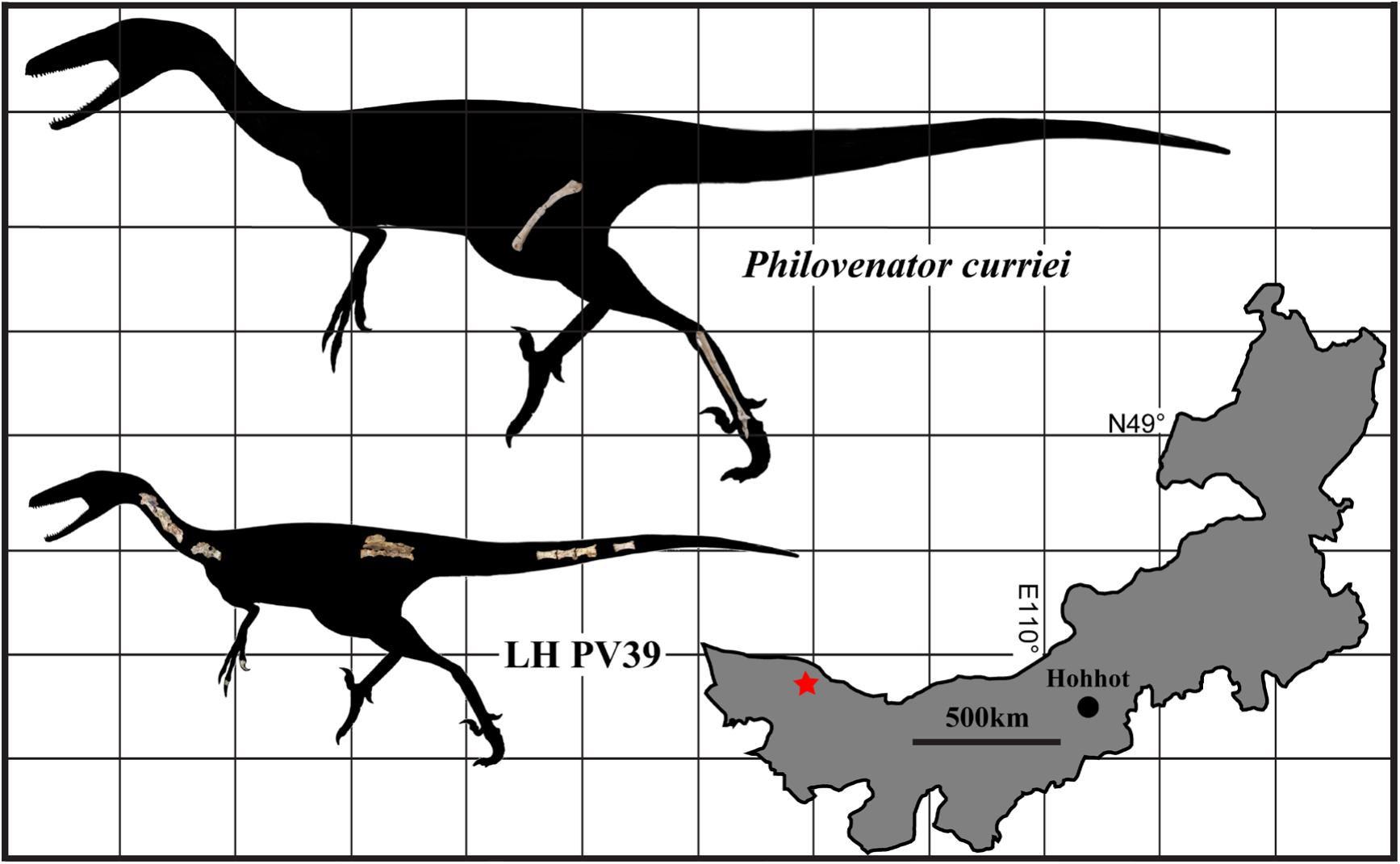
Skeletal silhouette with the representative elements showing LH PV39 (lower) is significantly smaller than *Philovenator* (IVPP V10597, upper) assuming the allometric growth pattern are identical in both taxa (*Almas* is about the same size of *Philovenator* given its femoral length is roughly as long as that of the latter); the simplified map showing the locality (red star) where LH PV39 was recovered. Each square=0.5m.

#### Cervicals

The exact positions of cervical vertebrae are difficult to determine given that relatively complete vertebral series are rarely known for troodontids and detailed vertebral descriptions are unavailable (Gao *et al*. 2012; Shen *et al*. 2017a; Shen *et al*. 2017b; Xu *et al*. 2017). The five relatively complete cervical vertebrae of LH PV39 are tentatively described as C4, C5, C6, C8 and C9, respectively (Fig. 2A-E), based on the increasingly greater separation of diapophyses and parapophyses, and the progressive reduction of offset between the anterior and posterior articular surfaces of the centrum posteriorly across the column. A partial left prezygapophysis in articulation with the diapophysis probably comes from C7 (data not shown). This element is only briefly described and will not be discussed further.

**Figure 2.**
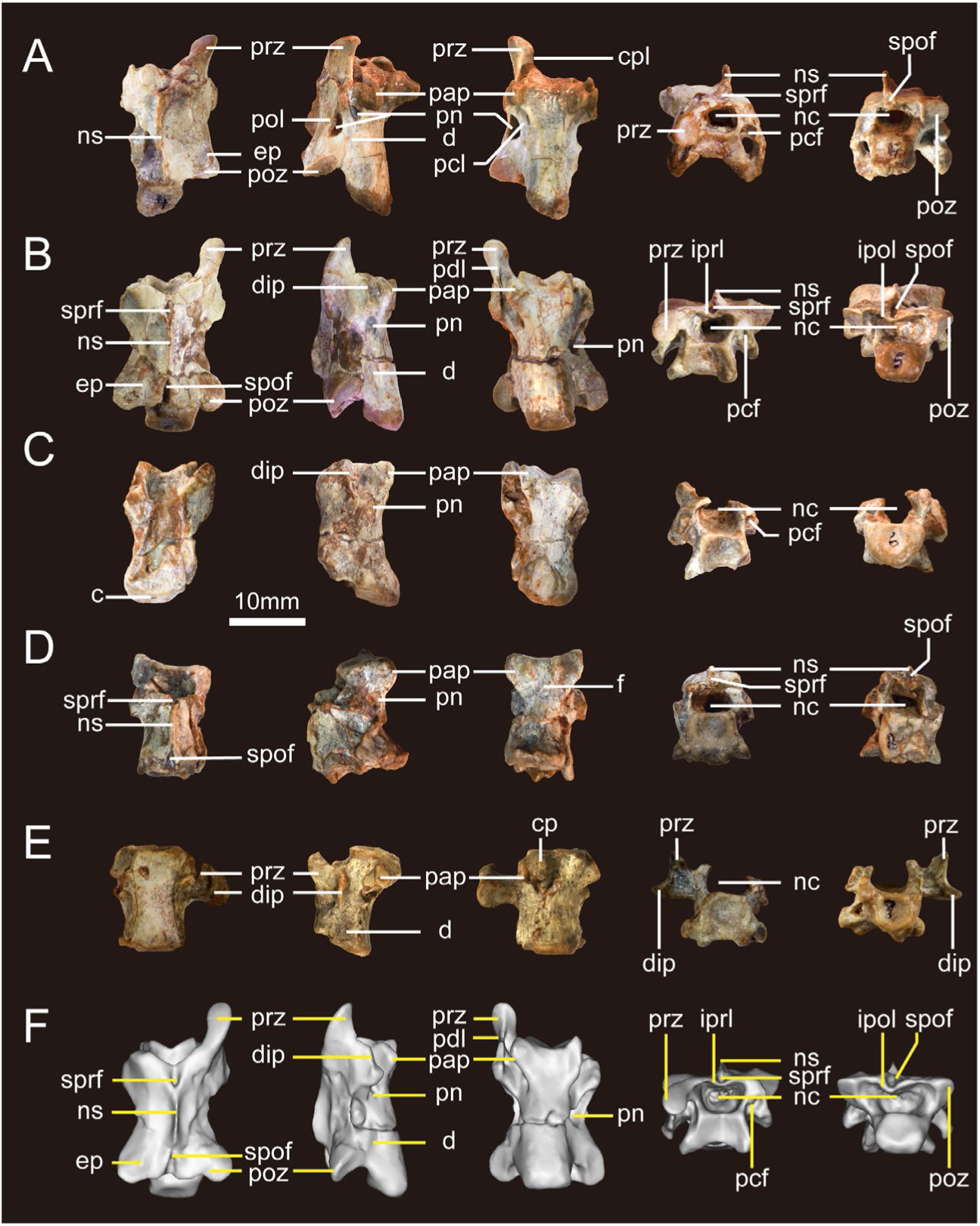
Photographs of cervical vertebrae of LH PV39. A-E, cervical 4 (A), 5 (B), 6 (C), 8 (D) and 9 (E) in dorsal, right lateral, ventral, anterior and posterior views from left to right columns, respectively. Abbreviations: c, centrum; cp, carotid process; cpl, centroprezygapophyseal lamina; d, depression; dip, diapophysis; ep, epipophysis; f, foramen; ipol, intrapostzygapophyseal lamina; iprl, intraprezygapophyseal lamina; nc, neural canal; ns, neural spine; pap, parapophysis; pcf, prezygapophyseal centrodiapophyseal fossa; pcl, posterior centrodiapophyseal lamina; pdl, prezygapodiapophyseal lamina; pn, pneumatic foramen; pol, postzygapodiapophyseal lamina; poz, postzygapophysis; prz, prezygapophysis; spof, spinopostzygapophyseal fossa; sprf, spinoprezygapophyseal fossa.

**Figure 3.**
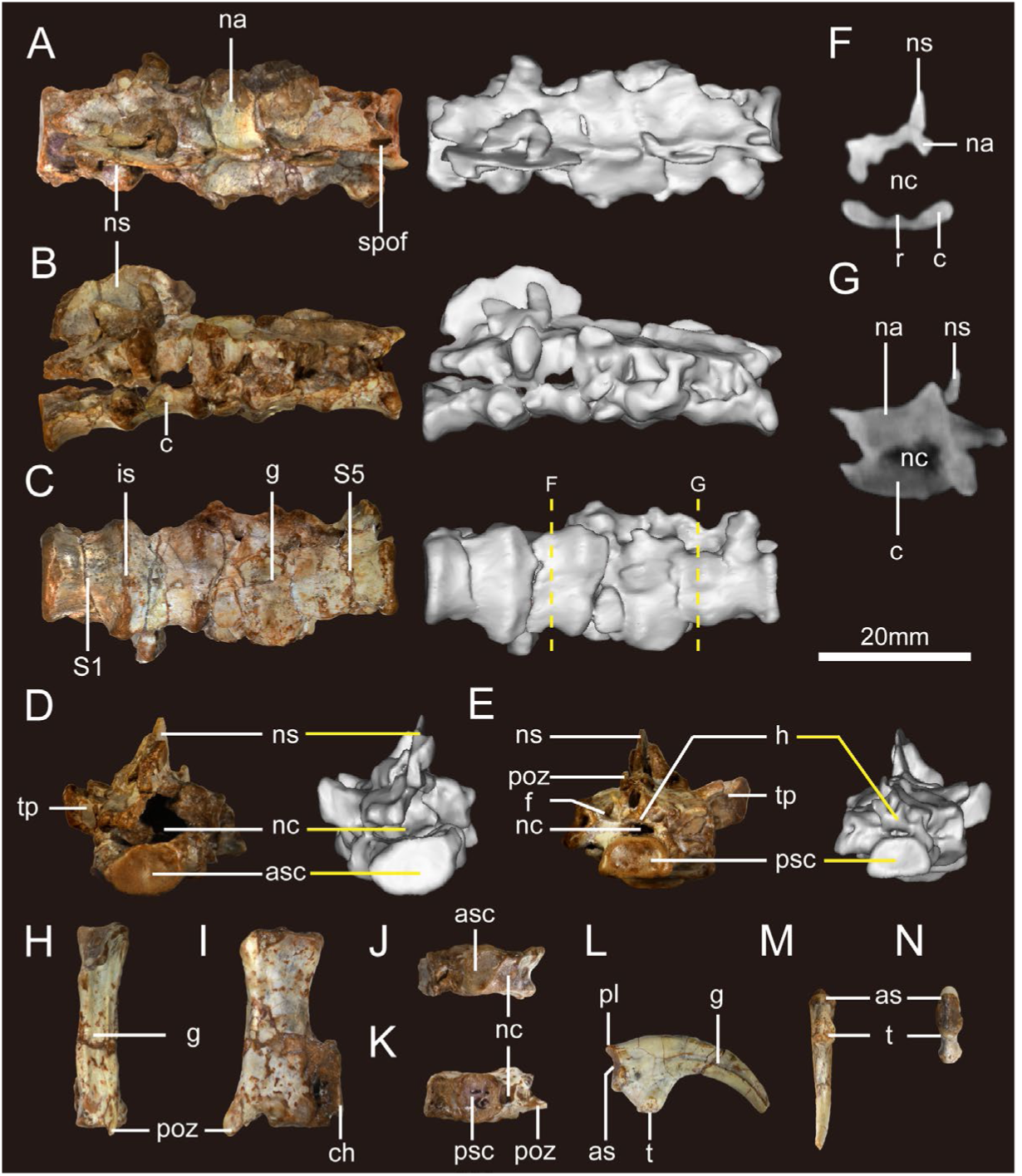
Photographs of synsacral, caudal vertebrae and manual ungual II-I of LH PV39. A-E, synsacral vertebrae (left) and the corresponding 3D reconstructions (right) in dorsal (A), right lateral (B), ventral (C), anterior (D) and posterior (E) views; the yellow dashed lines in C mark the positions of the slices shown in F and G; (F-G), coronal sections of sacral vertebra 2 (F) and 4 (G), showing the morphologies of the neural canal and centra; (H-K), CaB in dorsal (H), right lateral (I), anterior (J) and posterior (K) views; (L-N), manual ungual I-2 in right lateral (L), ventral (M) and proximal (N) views. Abbreviations: as, articular surface; asc, anterior articular surface of the centrum; c, centrum; ch, chevron; g, groove; h, hyposphenes; is, intercentral suture; na, neural spine; nc, neural canal; ns, neural spine; pl, proximodorsal lip; poz, postzygapophysis; psc, posterior articular surface of the centrum; r, midline ridge; S1-5, sacral vertebra 1-5, respectively; spof, spinopostzygapophyseal fossa; t, flexor tubercle; tp, transverse process.

All cervical vertebrae have a neural arch that is completely fused to the centrum without bearing any trace of an intervening suture (Fig. 2A-E). The centra of C4-C6 are each roughly 2 to 3 times longer than deep, whereas this value is less than 2 in C8 and C9, suggesting the anterior cervicals are more elongate than the posterior ones as in other paravians (Currie & Dong 2001; Pei *et al*. 2017a; Shen *et al*. 2017b; Xu *et al*. 2017). In fact, the peak centrum length is likely to be present at C5 as in other troodontids (Shen *et al*. 2017b; Xu *et al*. 2017), because cervicals both anterior and posterior to this point are much shorter (Table 1). It should be noted that the centra of C4 through C6 extend posteriorly beyond the postzygapophyses of the corresponding vertebra as in *Troodon* (Makovicky & Norell 2004) and *Sinornithoides* (Currie & Dong 2001), whereas it is slightly shorter than the postzygapophyses in more caudal cervicals (C8 and C9). The moderately concave anterior articular surface is wider than deep as in *Saurornithoides* (Norell & Hwang 2004) and *Mei* (Gao *et al*. 2012), in contrast to the relatively flat posterior surface (Fig. 2A-E). But they still represent amphiplatyan centra given the anterior surfaces are not as concave as that of the typical procoelous condition. In C4-C6, the anterior surface of the centrum is dorsally offset relative to the posterior surface as in other dinosaurs (Brochu 2003; Brusatte *et al*. 2012). This offset is progressively weaker in more posterior cervical vertebrae, suggesting the neck was held in a sMPC-Doid position in life. In addition, the anterior and posterior surfaces of the centrum of C4-C6 face anteroventrally and posterodorsally, respectively, when the centrum is horizontal, consistent with the interpretation of a curved neck.

**Table 1.** Measurements of vertebral elements of LH PV39.

|  | MLC | MHA | MWA | MHP | MWP | MLN |
| --- | --- | --- | --- | --- | --- | --- |
| C4 | 20.51 | 5.10 | 7.29 | 5.47 | 5.87 | 8.64 |
| C5 | 20.82 | 6.03 | 7.19 | 5.25 | 6.48 | 7.93 |
| C6 | 19.37 | 5.83 | 7.37 | 6.13 | 8.32 | - |
| C8 | 15.95 | 5.70 | 7.25 | 6.44 | 7.32 | 6.45 |
| C9 | 14.18 | 6.26 | 7.22 | 5.57 | 7.36 | - |
| S1 | 10.70 | 7.37 | 11.64 | - | 15.69 | - |
| S2 | 9.80 | - | 15.69 | - | 15.66 | - |
| S3 | 7.25 | - | 15.66 | - | 16.29 | - |
| S4 | 7.53 | - | 16.29 | - | 9.96 | - |
| S5 | 8.49 | - | 9.96 | 5.82 | 9.75 | - |
| CaA | 23.12 | 7.48 | 6.55 | 6.53 | 6.38 | - |
| CaB | 25.88 | 6.60 | 6.56 | - | 6.35 | - |
| CaC | 24.81 | 6.42 | 6.47 | 6.31 | 6.01 | - |
| CaD | 26.22 | 6.62 | 6.29 | 6.26 | 6.66 | - |

All values are in (mm). Abbreviations: C, cervicals; Ca, caudals; S, sacrals; MHA, maximum height of the anterior articular surface; MHP, maximum height of the posterior articular surface; MLC, maximum length of the centrum; MLN, maximum length of the neural spine; MWA, maximum width of the anterior articular surface; MWP, maximum width of the posterior articular surface.

In all cervicals, the parapophyses are located at the anteroventral corner on the lateral aspect of the centrum, unlike the condition in *Mei* in which the parapophyses are located as high as the dorsoventral midpoint of the centrum (Gao *et al*. 2012). The morphology and orientation of the parapophyses change along the neck as in dinosaurs generally (Brochu 2003; Brusatte *et al*. 2012; Madsen 1976). In C4∼C8, the parapophyses are triangular, plate-like structures in lateral view. In anterior view, they project lateroventrally in C4 (Fig. 2A), whereas they extend primarily laterally with their terminal ends located only slightly lower than the centrum in C8 (Fig. 2D). In C9, the parapophyses project entirely laterally, and they are significantly reduced in comparison to those of the anterior cervicals (Fig. 2E). In addition, the parapophyseal articular surfaces for the capitulum are strap-like and become progressively anteroposteriorly elongated posteriorly across C4 through C8. In contrast, they are ovoid and concave in C9 (Fig. 2E). The left parapophysis of C5 is excavated by a deep foramen (Fig. 2B), but it is likely to be a postmortem artifact as such a foramen is absent from the opposite side.

The lateral surface of each cervical centrum is moderately concave and excavated by a pneumatic foramen (*sensu* pleurocoel of some previous authors (Currie & Dong 2001; Gao *et al*. 2012)) located right above the base of the parapophysis, a feature that is also present in *Saurornithoides* (Norell & Hwang 2004) and *Mei* (Gao *et al*. 2012). However, the cervical vertebrae of *Sinornithoides* have two pneumatic foramina on each side (Currie & Dong 2001), with the anterior foramen located more posteriorly than the one present in LH PV39. The pneumatic foramen is slitlike in C4 through C6 (Fig. 2A-C), without presenting a shallow fossa surrounding it. However, it becomes rounded and more deeply excavated into the centrum posteriorly across the cervical series. An additional shallow depression is present posterior to the pneumatic foramen and close to the neurocentral junction (Fig. 2A, B). This shallow depression is more discernable in C4 and C5 than in the more posterior cervicals, is slightly deeper anteriorly, and becomes dorsoventrally expanded posteriorly giving it a triangular profile (Fig. 2A, B). In C4 and C5, this depression is even more elongated anterioposteriorly than the pneumatic foramen, whereas it becomes more oval, shallower and indistinct than the pneumatic foramen in C6, C8 and C9.

The ventral surfaces of the cervical centra are generally smooth, and there is no trace of a keel or hypapophysis, in contrast to the condition in *Troodon* (Makovicky & Norell 2004), *Saurornithoides* (Norell & Hwang 2004), *Mei* (Gao *et al*. 2012) and *Sinornithoides* (Russell & Dong 1993). In lateral view, the ventral surface is nearly flat in C4 and C5, whereas it is concave in more posterior cervicals (C6 through C9). The anterior portion of the ventral surface between the parapophyses bears a shallow triangular fossa in C4 and C5 (Fig. 2A, B), with one of the triangular vertex towards posteriorly. In C6 through C9, this fossa is laterally bound on each side by a moundlike carotid process (Fig. 2E), similar to the condition present in *Troodon* (Makovicky 1995), *Sinornithoides* (Russell & Dong 1993), *Mei* (Gao *et al*. 2012) and *Sinornithoides* (Currie & Dong 2001). Additionally, the carotid processes are more prominent in C9 than those present in C6 and C8 (Fig. 2E), and the distance between the two carotid processes increases posteriorly along the neck, unlike the condition in *Sinornithoides* in which the distance between the carotid processes decreases posteriorly across the cervical series (Currie & Dong 2001).

Though all known cervical centra are fused to their respective neural arches, relatively complete neural arches are only preserved in C4 and C5 (Fig. 2A, B). In dorsal view, the neural arch is transversely narrow at its midpoint, and the dorsal surface lateral to the neural spine is moderately concave (Fig. 2A, B). The neural spine of C4, C5 and C8 are preserved, but none of them are complete. Judging from the base, the neural spine is fairly thin and becomes shorter in the more posterior cervicals. From the base of the neural spine, a pair of spinoprezygapophyseal laminae extend anterolaterally and become confluent with the medial margin of the prezygapophysis on each side (Fig. 2B). Proximally, the spinoprezygapophyseal laminae demarcate the dorsolateral margin of the spinoprezygapophyseal fossa (*sensu* prespinal fossa (Brusatte *et al*. 2012)), which is located directly anterior to the neural spine. The spinoprezygapophyseal fossa is ventrally separated from the neural canal by the medially converged intraprezygapophyseal laminae, which are slightly ventrally protruded along the midline due largely to the postmortem deformation and give the neural canal a heart-shaped appearance (e.g. C5) (Fig. 2B). The neural canal is apparently wider than deep, and becomes progressively larger posteriorly along the neck.

Portions of the pre- and postzygapophyses are present on all cervicals. Although most of the left prezygapophysis has been eroded in C4 and C5, the right prezygapophyses are particularly well-preserved in these cervicals (Fig. 2A, B). In dorsal view, the prezygapophyses are slightly more widely spaced relative to the central midline than the postzygapophyses (Fig. 2A, B). The prezygapophyses extend primarily anterolaterally, and those in C5 are much slender and more anteriorly extended relative to those in C4. The prezygapophyseal articular surface is ovoid and smooth, with the long axis trending primarily anteroposteriorly (Fig. 2A, B). In lateral view, the articular surface is oriented anterodorsally, facing more anteriorly in C4 than that in C5, suggesting the neck would have been mostly curved at this position when the animal was alive.

In anterior view, the centroprezygapophyseal lamina extends medioventrally from the medial aspect of prezygapophysis, and finally contributes to the anterolateral margin of the neural canal (Fig. 2B). In addition to this lamina, the prezygapodiapophyseal lamina extends posteroventrally from the ventral aspect of the prezygapophysis and connects the diapophysis (Fig. 2A). This strut is transversely thick where it coalesces with the prezygapophysis, and becomes thinner as it extends posteroventrally, which not only forms the anterolateral margin of the large triangular flange of the diapophysis, but also contributes to the lateral margin of an anteriorly facing prezygapophyseal centrodiapophyseal fossa lateral to the neural canal (Brusatte *et al*. 2012; Wilson 1999) (Fig. 2A-C). This pocket is mostly prominent in C4 through C6, but whether it is also present in posterior cervicals remains unclear.

The articular surface of the diapophyses are strap-like with the long axis trending primarily anteroposteriorally in anterior cervicals before becoming ovoid in C7 (and more posterior cervicals?). In addition, this articular surface faces laterally as much as ventrally in C4 through C6, and closely approaches the parapophysis in anterior cervicals (Fig. 2A, B). In C9, it is located posterior to the parapophysis at a level that is roughly as high as the neurocentral suture while facing completely ventrally (Fig. 2E). This suggests the length, position, and orientation of the diapophysis changes posteriorly across the cervical series in LH PV39 as in other dinosaurs (Brochu 2003; Brusatte *et al*. 2012).

In addition to the prezygapodiapophyseal lamina, there are three more discrete laminae extending from the diapophysis in C4 through C6: two connecting with the centrum and one with the postzygapophysis. The anterior centrodiapophyseal lamina starts from the anteromedial corner of the diapophysis, and extends medially until reaching the neurocentral junction (Fig. 2A). This lamina demarcates the ventral margin of the prezygapophyseal centrodiapophyseal fossa which could be discerned in anterior view of C4 through C6 but is obscured by the pendant diapophysis when viewed laterally (Fig. 2A-C). The posterior centrodiapophyseal lamina, extends posteriorly from the posteromedial corner of the diapophysis, overhanging the pneumatic foramen above the parapophysis and gently coalesces with the centrum at approximately the anteroposterior midpoint of the centrum (Fig. 2A). This lamina is dorsoventrally broad where it starts, and becomes a thin osseous strut posteriorly. The postzygapodiapophyseal lamina links the diapophysis with the postzygapophysis. This lamina, along with the prezygapodiapophyseal lamina, defines the lateral margins of the large triangular diapophysis in all cervicals. Posterior to the diapophysis and on the lateral aspect of the neural arch, there is a deeply excavated pneumatic foramen dorsally and ventrally demarcated by the postzygapodiapophyseal and the posterior centrodiapophyseal laminae, respectively (Fig. 2A, B), a feature that is also present in most other troodontids (Norell & Hwang 2004; Shen *et al*. 2017b) but apparently absent in *Mei* (Gao *et al*. 2012). CT images show that this foramen does not deeply excavate the neural arch in C4 (data not shown). In C5 and C6, this fossa invades the posterior aspect of the diapophysis (Fig. 2B, C).

Complete paired postzygapophyses are known only in C5 (Fig. 2B), and the right postzygapophysis of C4 preserves some morphology (Fig. 2A). In dorsal view, the postzygapophyses do not extend as posteriorly as the corresponding centrum in C4 and C5, differing from the condition in C8. The postzygapophyseal articular surface is ovoid and moderately concave in C4 (Fig. 2A), in contrast to the fairly flat and transversely narrow condition in C5 (Fig. 2B). In addition, this surface faces primarily posteroventrally and only slightly laterally in C4, whereas it faces posteroventrally as much as laterally in C5. Although the postzygapophyses were completely eroded in C9, there is a smooth, transversely wide surface present on the left margin of the posterior wall of the neural canal, which extends to the lateral surface of the neural arch. However, it is unlikely to be an articular surface given a similar structure is neither present in the opposite side of this vertebra nor in other cervical vertebrae, therefore it could be either a pathological structure or merely a preservational artifact.

Unlike the prezygapophyses, the postzygapophyses do not connect with the main body of the neural arch through a bony neck (Fig. 2A, B). In dorsal view, the postzygapodiapophyseal and intrapostzygapophyseal laminae contribute to the lateral and posterior margins of the postzygapophyses, respectively (Fig. 2A, B). In posterior view, the intrapostzygapophyseal lamina from both sides converge along the midline, where they protrude slightly ventrally due primarily to postmortem deformation, giving the neural canal a heart-shaped appearance (Fig. 2B). Here, the intrapostzygapophyseal laminae also form the bottom of a well-developed spinopostzygapophyseal fossa (or postspinal fossa (Wilson *et al*. 2011)) (Fig. 2A, B, D). The neural spine and the postzygapophysis are connected by the spinopostzygapophyseal lamina which is dorsally raised as a slight lip that contributes to the medial margin of the postzygapophysis (Fig. 2B). In C4, the spinopostzygapophyseal fossa is more prominent and anteroposteriorly elongate than the spinoprezygapophyseal fossa. However, in C5, the spinoprezygapophyseal fossa is deeply excavated into the neural spine, and roughly as long as the spinopostzygapophyseal fossa. Incomplete preservation of spinopre- and spinopostzygapophyseal fossae in C8 precludes further comparisons.

The dorsal surface of the postzygapophysis is not flat because an epipophysis is present as in *Mei* (Gao *et al*. 2012) and *Saurornithoides* (Norell & Hwang 2004), perhaps indicating relatively strong neck musculature in LH PV39. In C5, the epipophysis is located at the center of the postzygapophysis when viewed dorsally (Fig. 2B), whereas it is located much closer to the postzygapophyseal posterolateral margin in C4 (Fig. 2A), suggesting the position of the epipophysis is various across the neck. In C4, there is a weak ridge extending anteriorly from the epipophysis for a distance that is roughly as long as the epipophysis itself, but it does not form a lamina anteriorly connecting to the prezygapophysis as in *Alioramus* (Brusatte *et al*. 2012).

The available cervical series of LH PV39 also includes a partial left prezygapophysis in association with diapophysis (data not shown). Because the prezygapophysis and the diapophysis are closely situated, it is probably coming from a cervical posterior to C6. The articular surface of this prezygapophysis is ovoid, fairly flat and slightly more expanded than that in C9, and its base is also anteroposteriorly longer than that of C9. Therefore, this prezygapophysis is most likely coming from C7 given that the bases of the prezygapophyses are preserved in both C8 and C9.

#### Synsacrum

The synsacrum consists of five co-ossified vertebrae, though the intercentral sutures are still visible (Fig. 3A-E). Among troodontids, Late Cretaceous taxa such as *Saurornithoides* (Norell *et al*. 2009), *Troodon* (Norell *et al*. 2009), *Gobivenator* (Tsuihiji *et al*. 2014), *Latenivenatrix* (van der Reest & Currie 2017), *Zanabazar* (Norell *et al*. 2009) and an unnamed specimen collected from Two Medicine Formation (van der Reest & Currie 2017) each has a sacrum consisting of six vertebrae, whereas there are only five sacral vertebrae in the Jurassic and Early Cretaceous taxa *Mei* (Gao *et al*. 2012), *Sinusonasus* (Xu & Wang 2004b), *Daliansaurus* (Shen *et al*. 2017a), *Archiornis* (Pei *et al*. 2017a) and *Sinovenator* (Xu *et al*. 2002) as in many other paravians (Godefroit *et al*. 2013; Norell & Makovicky 1997; Ostrom 1976; Turner *et al*. 2012; Xu *et al*. 2011b). Character optimization indicates that five sacral vertebrae is the plesiomorphic condition for troodontids and the addition of a sixth sacral vertebrae occurred one or more times during the evolution of troodontids.

All sacral centra are well preserved. In ventral view, each sacral centrum is constricted at its midpoint (Fig. 3A-E), but CT images show that they are not spool-like elements thus unlike the condition in most other theropods (Brochu 2003; Brusatte *et al*. 2012; Norell *et al*. 2009; van der Reest & Currie 2017; Zanno *et al*. 2011), instead they are dorsoventrally compressed as in *Sinovenator* (Xu *et al*. 2002), dromaeosaurids *Rahonavis* (Forster *et al*. 1998), *Buitreraptor* (Novas *et al*. 2018), *Mahakala* (Turner *et al*. 2007; Turner *et al*. 2011) and the oviraptorosaurs *Microvenator* (Makovicky & Sues 1998) and *Chirostenotes* (Currie & Russell 1988; Sues 1997) (Fig. 3A-G). In LH PV39, a broken surface reveals that the centrum of S2 is a plate-like element with its dorsal aspect that is excavated by shallow depressions, suggesting that this region has been substantially eroded. But CT images revealed the centra of the non-eroded S4 and S5 are also compressed, demonstrating the compressed nature of the sacral centra might be one of the distinct features for LH PV39 rather than a result of postmorteum deformation or erosion. The anteroposterior length of individual centra decreases posteriorly across S1 through S3, reaching a minimum length of 7.25 mm in S3 before increasing again posteriorly (Table 1). It should be noted that the length variation of the sacral vertebrae is extensive among paravians, and LH PV39 makes a noteworthy contribution to this morphological diversity. In most paravians, the sacral vertebrae are subequal in length (e.g. *Anchiornis* (Pei *et al*. 2017a), *Mei* (Xu & Norell 2004), *Sinovenator* (Xu *et al*. 2002), *Buitreraptor* (Novas *et al*. 2018) and *Zanabazar* (Norell *et al*. 2009)) as in most theropod dinosaurs. The sacral vertebrae of *Mahakala* are similar in length except for the first and last vertebrae, which are significantly longer than the middle ones (Turner *et al*. 2011). Therefore, LH PV39 resembles *Mahakala* in both the compressed morphology and the length variation of sacral centra.

Although the vertebrae are completely fused, the intercentral contacts remain visible as the contact regions are more expanded than the midlength of the centrum (Fig. 2). The transverse widths of the centra increase posteriorly in S1 through S3, reaching a maximum width of 16.29 mm in S3 and then decrease again posteriorly as in *Sinovenator* (Xu *et al*. 2002), *Microraptor* (Xu *et al*. 2000), and *Archiornis* (Pei *et al*. 2017a). Despite the intercentral junctions of S1 and S2, and S2 and S3 being of similar width, the anterior articular surface of the first sacral centrum is transversely narrower than the posterior surface of the corresponding centrum, whereas the anterior surface of the fourth is transversely wider than the posterior surface of the corresponding cnetrum (Fig. 3C). These suggest the sacral central width increase and decrease dramatically in S1 and S4, respectively. The last centrum (S5) is the narrowest one among the sacrals (Fig. 3C).

The ventral surface of the centrum in S1 is smooth and gently convex, without any sign of keels and grooves as in other theropod dinosaurs (Brusatte *et al*. 2012; Wang *et al*. 2016). However, a shallow groove extends posteriorly from the junction of S1 and S2 until the midlength of S5, cutting through the expanded intercentral junctions between S2 and S3, S3 and S4, and S4 and S5 (Fig. 3C). The transverse width of this groove varies little as it extends, which approximately equals to one third the width of the last sacral. A similar midline groove is also present in *Latenivenatrix* (van der Reest & Currie 2017), *Mahakala* (Turner *et al*. 2011), *Balaur* (Brusatte 2013), *Velociraptor* (Brusatte 2013) and *Bambiraptor* (Burnham 2004), but is absent in *Saurornitholestes* (Norell *et al*. 2009; van der Reest & Currie 2017), *Archiornis* (Pei *et al*. 2017a) and *Talos* (Zanno *et al*. 2011), among other paravians. In *Zanabazar*, the ventral midline groove can be discerned only around the intercentral junctions (Norell & Hwang 2004). The extent of this groove is also variable among paravians. It marks only the second through the fourth sacrals in *Balaur* and *Velociraptor* (Brusatte 2013), whereas it starts from the posterior half of the second sacral in *Mahakala* (Turner *et al*. 2011). In *Latenivenatrix* (van der Reest & Currie 2017), this groove is present in S3 through S5, whereas it is present only on the third sacral in *Bambiraptor* (Burnham 2004). Unlike other theropod dinosaurs in which the ventral surfaces of centra are marked by a series of anteroposteriorly trending striations adjacent to the intercental contacts (Brusatte *et al*. 2012), these regions are fairly smooth in LH PV39.

Articular surfaces of the sacral centra are mostly invisible, and even CT images could not reveal details of the intercentral junctions, confirming that the sacrals are completely fused. Therefore, only the anterior and posterior intercentral surfaces of S1 and S5 are exposed, respectively (Figs. 3D, E). The anterior surface of S1 is ovoid, nearly flat, being approximately twice as transverse wide as depth similar to that seen in *Buitreraptor* (Novas *et al*. 2018) (Fig. 3D). In lateral view, this surface is almost straight vertically, which is slightly ventrally offset relative to the intercentral junction of S1 and S2 (Fig. 3B). In contrast, the trapezoid posterior surface of S5 is concave, with the straight ventral margin that is transversely wider than the dorsal margin (Fig. 3E), reminiscent of that in *Latenivenatrix* (van der Reest & Currie 2017). In lateral view, this surface faces slightly dorsally, and extends posteriorly as far as the postzygapophysis (Fig. 3B). In addition, the posterior surface of S5 is slightly ventrally offset relative to the intercentral junction of S4 and S5 as in other theropod dinosaurs (Brusatte *et al*. 2012).

The neural arch is fused to the centrum in all sacrals, and the neural spines are also fused into a single apron as in other theropods (Barsbold 1974; Barsbold *et al*. 2000; Brochu 2003; Xu *et al*. 2006) (Fig. 3A). Anteriorly, a pair of spinoprezygapophyseal laminae extend from the neural spine and demarcate a spinoprezygapophyseal fossa in S1, but little can be said about the prezygapophyses themselves because of serious erosion. The zygapophyses of the remaining sacrals are reduced and likely to have been fused. In dorsal view, the dorsal surface of the neural arch lateral to the neural spine is moderately concave, but the lateral surfaces of the neural spine are fairly flat (Figs. 3A-C). The neural arches are transversely expanded at the anteroposterior midpoint, where a pair of transverse processes project laterally (Figs. 3A, C). The right transverse process of S2 is preserved, it extends slightly dorsolaterally while terminates in a dorsoventrally expansion for the sacral rib (Fig. 3B). No sacral ribs are preserved in association with the sacrum, suggesting the ribs and sacral vertebrae were not fused.

Only the left postzygapophysis is well preserved in S5 (Fig. 3E). It projects posterodorsally and terminates roughly at a same level of the posterior surface of the centrum, as has been mentioned above. The postzygapophyseal articular surface is fairly small, which faces more laterally than ventrally (Fig. 3E). Lateral to the base of the postzygapophysis, the neural arch is excavated by a deep, posterodorsal facing foramen on each side such that the posterior margin of the neural arch between the transverse process and the postzygapophysis is demarcated by a deep V-shaped notch when viewed dorsally (Fig. 3A). The medial margin of the postzygapophysis connects with the neural spine proximally via the spinopostzygapophyseal lamina, and the left and right spinopostzygapophyseal laminae together demarcate a deep and elongate spinopostzygapophyseal fossa (Fig. 3A). The presence of both spinoprezygapophyseal and spinopostzygapophyseal fossae suggest five sacral vertebrae seems to be a reasonable estimate for the synsacrum of LH PV39. Ventral to the postzygapophysis and on the posterior surface of the neural arch, there is a pair of very small posteriorly facing articular surfaces, presumably representing the hyposphenes (Fig. 3E). The surface between the hyposphenes is flat and faces posteroventrally. Immediately ventral to the possible hyposphenes, the extremely dorsoventrally compressed neural canal is visible, with a transverse width of 6.25 mm and a depth of 1.65 mm (Fig. 3E). The dorsoventrally compressed neural canal is another peculiar character of LH PV39, in addition to the compressed sacral vertebrae.

#### Caudals

Four middle caudal are preserved in sequence. Comparisons with other troodontids preserving nearly complete caudal series suggest they are likely to fall somewhere within the range of Ca10-20 (Currie & Dong 2001; Gao *et al*. 2012; Shen *et al*. 2017a; Shen *et al*. 2017b; Tsuihiji *et al*. 2014; Xu *et al*. 2017; Xu *et al*. 2002), but their exact positions are difficult to determine. Therefore, they are referred to as CaA, CaB, CaC and CaD, respectively, from anterior to posterior.

The centra are low, and roughly three times longer than deep (Figs. 3H-K; Table 1). In addition, the transverse processes are reduced to weakly developed nubbins present on the lateral surfaces such that they are difficult to discern. The neural arch and centrum are clearly completely fused in all known caudals, and there is no sign of a neurocentral suture between the two components (Figs. 3H-K). The centra are rod-like and amphiplatyan, and all intercentral articular surfaces are only slightly concave. In lateral view, there is no offset between the anterior and posterior articular surfaces, but the dorsal and ventral margins of the vertebra are gently concave at the midpoint as in most other dinosaurs (Averianov & Sues 2016; Norell *et al*. 2009) (Fig. 3H). The lateral surfaces apparently lack pneumatic foramina, though the rough surfaces suggest the caudals have been seriously weathered. The ventral surface of the centra are marked by a distinct shallow groove as in other troodontids (Averianov & Sues 2016; Zanno *et al*. 2011). The posterior articular surface extends slightly to the ventral aspect of the vertebrae, representing the facet for chevron articulation.

Neural arches are present in all preserved caudal vertebrae, which are only slightly longer than the corresponding centrum (Fig. 3H), unlike the condition in dromaeosaurids in which the elongate prezygapophyses are usually several times longer than the centrum (Norell & Makovicky 2004; Ostrom 1969). A neural spine is absent from all vertebrae, and its position is taken by an anteroposteriorly extended groove as in all known troodontids (Currie & Dong 2001; Gao *et al*. 2012; Norell & Hwang 2004; Norell *et al*. 2009; Norell *et al*. 2000; Russel 1969; Russell & Dong 1993; Shen *et al*. 2017a; Tsuihiji *et al*. 2014; Xu *et al*. 2002; Xu & Wang 2004b; Zanno *et al*. 2011) (Fig. 3f), which is deeper than the one present on the ventral aspect of the centrum. The zygapophyses are set at a low angle relative to the centrum (Figs. 3H-I). No caudal vertebra is preserved with complete prezygapophyses, but the prezygapophyses are slightly more widely positioned than the postzygapophyses relative to the midline judging from their bases (Fig. 3I). Between the prezygapophyses, there is a deeply excavated, well defined anteriorly facing spinoprezygapopgyseal fossa, which is roofed by the posteromedially converged intraprezygapophyseal laminae. A distinct sharp ridge starts from the medial aspect of the prezygapophysis, extending posteriorly and becoming confluent with the dorsal margin of the postzygapophyses posteriorly (Fig. 3H). These ridges define the lateral extent of the groove on the dorsal surface of the neural arch. In addition to this ridge, another ridge extends from the dorsolateral aspect of the prezygapophysis and continues with the postzygapophyses separating the lateral surface of the neural arch from its dorsal surface. The postzygapophysis on the right side of CaA and CaB, and that on the left side of CaC are preserved, with their articular surfaces extending only slightly posterior to the centrum (Fig. 3H-I). The articular surfaces are fairly small and flat, tapering distally and facing completely laterally as in *Zanabazar* (Norell *et al*. 2009). This suggests the prezygapophyseal articular surfaces of the succeeding caudal are fairly short and face primarily medially, and the lateral bending capability within the caudal column was rather restricted for LH PV39. Other detailed morphology of the postzygapophysis remains unclear due to erosion. No trace of a spinopostzygapophyseal fossa is present, a feature that differs from *Urbacodon* (Averianov & Sues 2016). In anterior or posterior views, the neural canal is oval shaped, which is roughly as big as the spinoprezygapophyseal fossa (Fig. 3J-K), unlike the condition in *Urbacodon* in which both the spinopre- and spinopostzygapophyseal fossae are larger than the neural canal (Averianov & Sues 2016).

The preserved fragmentary chevrons in association with CaA and CaB are flattened plate-like elements (Fig. 3I). The ventral surface of the chevron bears a midline groove that is shallow and transverse broad. Other morphologies are difficult to describe due to poor preservation.

#### Manual Unguals

Only two unguals were recovered, both are strongly mediolaterally compressed and recurved (Figs. 3L-N). The side and digit that each ungual belongs to is difficult to determine, but both are identified as manual unguals. One nearly complete ungual is elongate and sharply pointed distally. It is 24.2 mm along the outer curvature, missing only a very small portion of the distal tip. Proximally, the ovoid articular surface is 5.8 mm deep and 2.3 mm wide, which is divided into two nearly equal concave facets by a prominent vertical medial ridge (Fig. 3N). In lateral view, the dorsalmost point of the proximal articular surface contributes to a proximodorsal lip that projects primarily posteriorly and only slightly dorsally (Fig. 3L). The well-developed flexor tubercle is ventrally rugose (Figs. 3M-N) and is separated from the ventral margin of the proximal articular surface by a dorsoventrally broad transverse groove that is not as deep as those seen in other maniraptorans (Bell *et al*. 2015; Novas *et al*. 2005). A distinct groove is present on both sides for vascular supply to the keratinous sheath. The groove on the left side extends from the distal extremity of the element, and becomes shallower as it extends proximally to a position between the flexor tubercle and the proximal articular surface. Although the groove on the right side is roughly as deep as the medial one, it proximally terminates anterior to the flexor tubercle. This ungual is interpreted as a manual ungual based on the following criteria: (1) curvature of the ungual is approximately 120 degrees (following the method of ref. (Pike & Maitland 2004)); (2) the proximal articular surface is taller than its transverse width; (3) the presence of a proximodorsal lip; (4) the presence of a transverse groove separating the proximal articular surface from the well-developed flexor tubercle; and (5), the proximal articular surface comprises two nearly equal facets. The other preserved ungual provides insufficient morphology to merit description as most of its distal and proximal ends have been broken off, but can be identified as a manual ungual based on its comparable size to the previously described ungual.

## Phylogenetic Analyses

LH PV39 is referable to Troodontidae based on the following combination of features: presence of cervical epipophyses above the postzygapophyseal facets, anterior cervicals centra extend beyond the posterior limit of the corresponding neural arch, fused sacral zygapopyses, and the presence of a midline groove on the dorsal aspect of caudal neural arch.

To test the phylogenetic position of LH PV39, we conducted several phylogenetic analyses by coding LH PV39 in different strategies into the modified published data matrices designed to analyze coelurosaurian relationships. We first scored LH PV39 as a separate OTU, which are designed to explore the possible effects of including LH PV39 morphological data on the interrelationships of troodontids and test the phylogenetic position of LH PV39. Because LH PV39 was recovered from a locality that is fairly geographically close to that of *Philovenator*, and size comparisons of LH PV39 and *Philovenator* do not preclude the referral of LH PV39 to the latter taxon, we then coded the remains of LH PV39 and *Philovenator* as a composite OTU in additional analyses, to test whether the referral of LH PV39 to *Philovenator* alters the troodontine affinities of that taxon. Each of these coding strategies were performed using the matrix that was modified from Senter et al. (2012) and Turner et al (2012), respectively, to verify the phylogenetic position of LH PV39.

### Analyses using modified data matrices of Senter et al. (2012)

For the data matrices modified from Senter et al. (2012) (see Supplementary Information 1), three new characters (characters 395, 396 and 397) were added, and these are present below:

395: sacral centra spool-like (0) or dorsoventrally compressed and plate-like (1) (newly added character).

396: sacral neural canal oval (0) or dorsoventrally compressed (1) (newly added character).

397: sacral: centra similar in width (0), or middle centra narrower than the first and the last centra (1), middle centra wider than the first and the last centra (2), centra width constantly decrease posteriorly (3), centra width constantly increase posteriorly (4).

In addition, the matrices were expanded by adding *Linheventator* (Xu *et al*. 2011a), *Philovenator* (Xu *et al*. 2012), *Almas* (Pei *et al*. 2017b), and LH PV39 (in Analysis 1), making the final matrices consist of 113 and 112 taxa across 397 characters for Analysis 1 and 2, respectively (see Supplementary Information). The data matrices were analyzed with equally weighted parsimony using the software package of TNT v.1.5 (Goloboff et al. 2008). The traditional tree search strategy was conducted performing 1000 replicates of Wagner trees. This tree searching strategy aims to obtain all the most parsimonious resolutions.

When coding LH PV39 as a separate OTU, the analysis yielded 174 most parsimonious trees, each with a length of 1373 steps. The strict consensus has a consistency index of 0.361 and a retention index of 0.803 (Fig. 4). The interrelationships of Troodontidae are not well-resolved. Bootstrap and Bremer support values for nodes within Troodontidae are low in the strict consensus of these trees (Fig. 4), and LH PV39 is resolved in a polytomy of troodontids outside Troodontinae (Fig. 4). Among the known 174 parsimonious trees, the position of LH PV39 oscillates between *Anchiornis* and the common ancestor of *Linhevenator* and *Troodon* (Fig. 4). In addition, this analysis yielded *Almas* as a sister taxon of *Linhevenator* and *Philovenator*.

**Figure 4.**
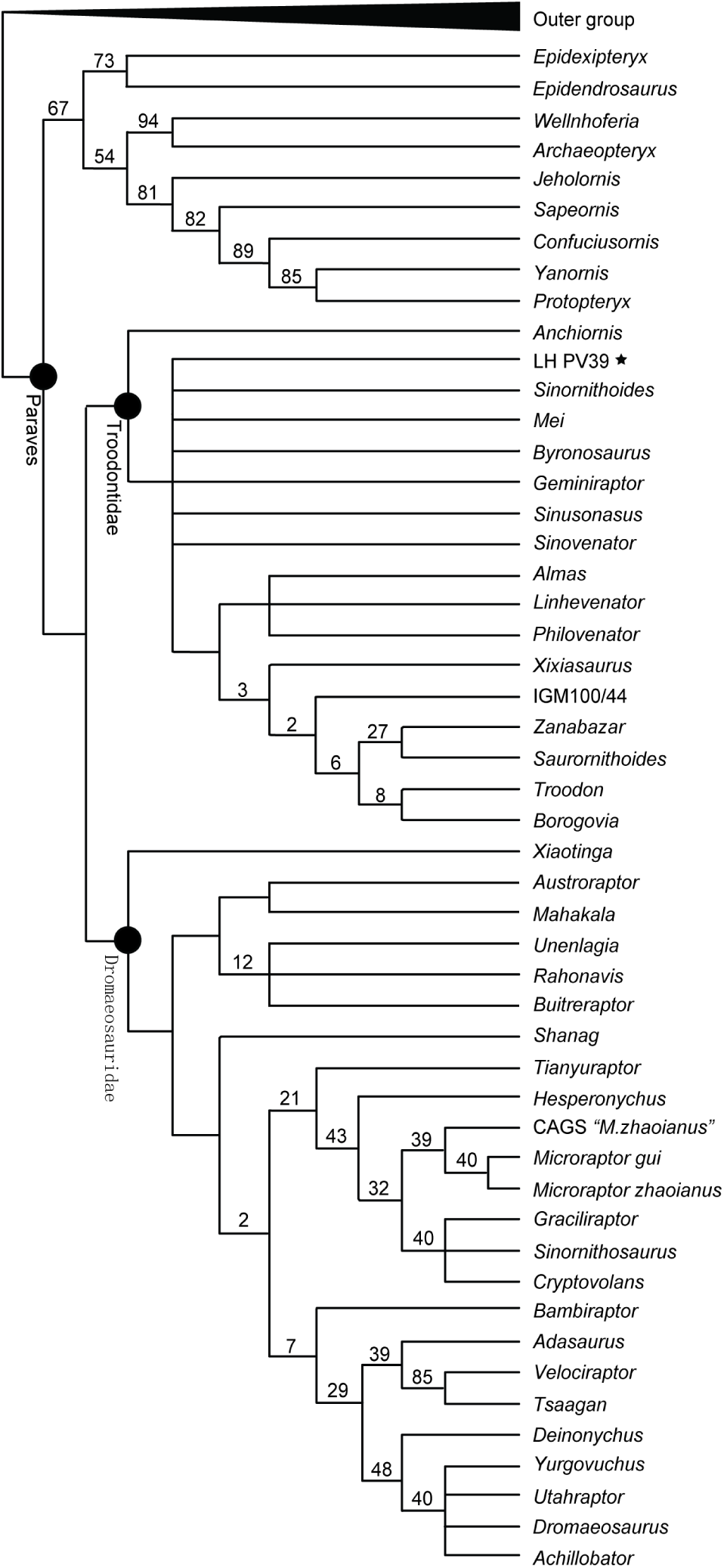
Simplified strict consensus cladogram showing the Paraves interrelationships when coding LH PV39 as a separate OTU using modified data matrix of Senter et al. (2012). The phylogenetic analysis resultant in 174 most parsimonious trees of 1373 steps, each with a consistency index of 0.361 and a retention index of 0.803. Values above nodes represent bootstrap percentages (%), and values lower than 20% are not shown.

When coding LH PV39 and *Philovenator* as a composite OTU, the analysis resulted in 93 most parsimonious trees and each with a length of 1374 steps. The strict consensus has a consistency index of 0.360 and a retention index of 0.803 (Fig. 5). The Bootstrap support values increase relative to those of the previous analysis. The interrelationships of Troodontidae are basically well resolved, the phylogenetic position of *Almas* has nothing different from that in the previous analysis, and the composite OTU of *Philovenator* and LH PV39 is unrefutably recovered as the sister taxon of *Almas* and *Linhevenator* (Fig. 5). This suggests that the topology of the strict consensus was not affected when incorporating LH PV39 into *Philovenator*.

**Figure 5.**
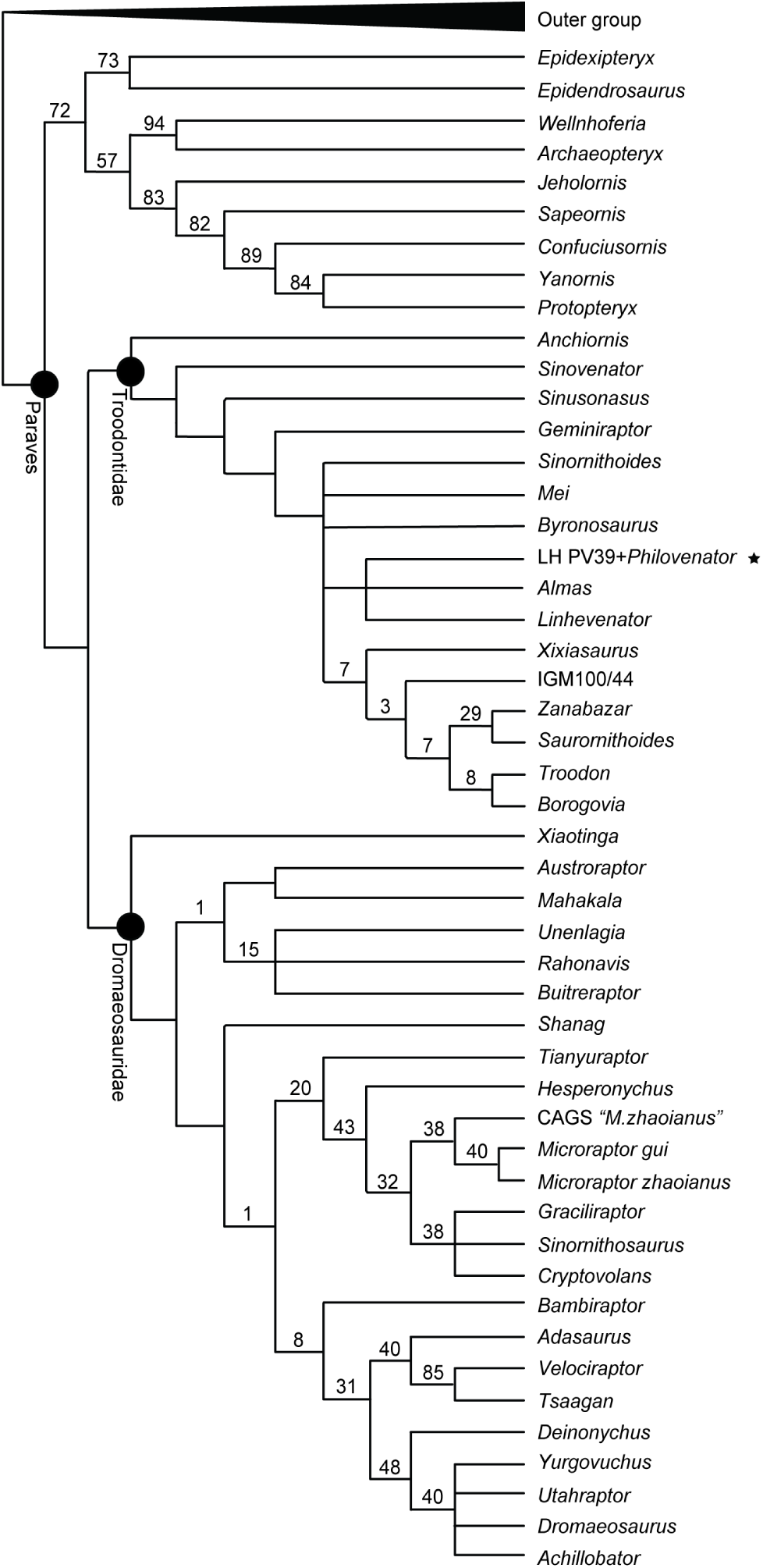
Simplified strict consensus cladogram showing the Paraves interrelationships when coding LH PV39 and *Philovenator* as a composite OTU using modified data matrix of Senter et al. (2012). The phylogenetic analysis resultant in 93 most parsimonious trees of 1374 steps, each with a consistency index of 0.361 and a retention index of 0.803. Values above nodes represent bootstrap percentages (%), and values lower than 20% are not shown.

### Analyses using modified data matrices of Turner et al. (2012)

For data matrices modified from (Turner *et al*. 2012), the three newly added characters 478, 479 and 480 are identical to the characters 395, 396 and 397 added to Senter et al. (2012), and the additional modifications of the character states of 57 are present below: 57:

> Paroccipital process: straight, projects laterally or posterolaterally (0), or distal end curves ventrally, pendant (1), or distal end curves dorsally (2) (newly added state).

A total of 75 character states of *Almas* (MPC-D 100/1323) in the original published data matrix has been updated according to the detailed osteological description of this specimen (Pei *et al*. 2017b) (see Supplementary Information). By adding *Linheventator* (Xu *et al*. 2011a), *Philovenator* (Xu *et al*. 2012), and LH PV39 (in Analysis 3), the final matrices yielded 120 and 119 taxa across 480 characters, respectively, depending on whether LH PV39 and *Philovenator* were coded as a composite or separate OTU (see Supplementary Information). Characters 6, 50, 52 and 147 were excluded in the analyses with 50 characters treated as ordered, in consist with the original analyses (Turner *et al*. 2012).

The data matrices were analyzed with equally weighted parsimony using the software package of TNT v.1.5 (Goloboff et al. 2008). The traditional tree search strategy was conducted performing 99999 replicates of RAM trees followed by TBR branch swapping (holding 20 trees per replicate), with the exclusion of *Unenlagia*, *Pedopenna*, *Epidendrosaurus* The best trees obtained at the end of the replicates were subjected to a final round of TBR branch swapping, and zero-length branches were collapsed if they lacked support under any of the most parsimonious reconstructions. To further improve the resolution, *Almas*, *Byronosaurus*, *Xixiasaurus*, *Pyroraptor*, *Hesperonychus* were reduced from the consensus, resulting in 112 and 128 parsimonious trees when LH PV39 was coded in different strategies. This tree searching strategy aims to obtain all the most parsimonious resolutions.

When coding LH PV39 as a separate OTU, the analysis yielded 112 most parsimonious trees, each with a length of 2089 steps. The Bootstrap and Bremer support values for are significantly low, and the consistency and retention indices of the strict consensus are confident. Almost all major branches are well-resolved and LH PV39 is recovered as the sister taxon of *Mei* and *Sinovenator* (Fig. 6). When coding LH PV39 and *Philovenator* as a composite OTU, the analysis produced 128 most parsimonious trees, each with a length of 2090 steps (Fig. 7). Similarly, the Bootstrap and Bremer support values are not ideal, and the consistency and retention indices of the strict consensus are not confident though the resolution of the coelurosaurian interrelationships is better than the result of when LH PV39 was coded as a separated OTU (Fig. 7). The composite OTU of *Philovenator* and LH PV39 is resolved as the sister taxon of *Linhevenator*, suggesting the referral of LH PV39 to *Philovenator* does not affect the topology of the consensus nor the relationship between *Philoventor* and *Linhevenator*.

**Figure 6.**
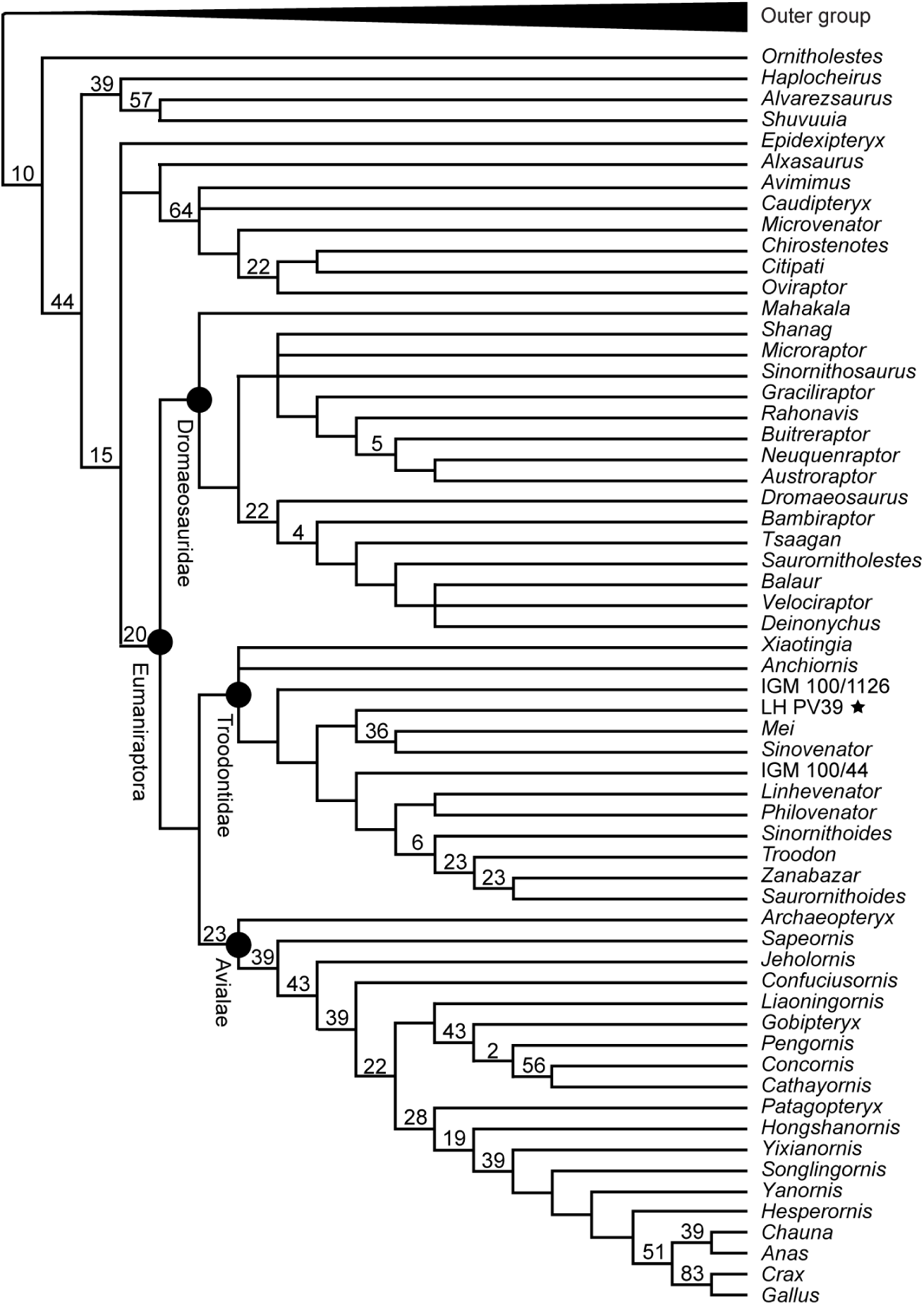
Simplified strict consensus cladogram showing the Paraves interrelationships when coding LH PV39 as a separate OTU using modified data matrix of Turner et al. (2012). The phylogenetic analysis resultant in 112 most parsimonious trees of 2089 steps.

**Figure 7.**
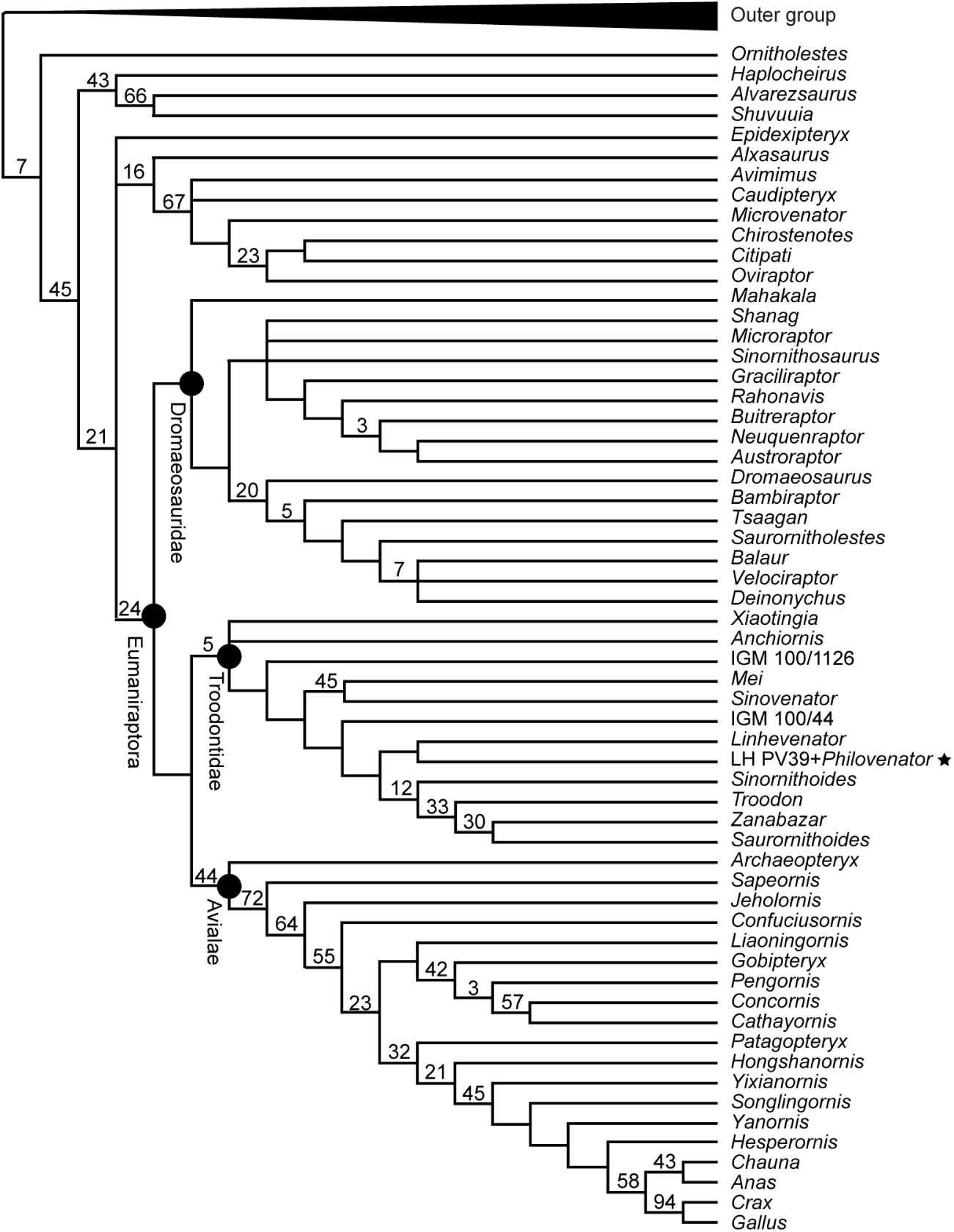
Simplified strict consensus cladogram showing the Paraves interrelationships when coding LH PV39 and *Philovenator* as a composite OTU using modified data matrix of Turner et al. (2012). The phylogenetic analysis resultant in 128 most parsimonious trees of 2090 steps.

## Discussion

Among troodontids, LH PV39 differs from *Mei* in the absence of keel in anterior cervicals (Gao *et al*. 2012), differs from *Sinornithoides* and *Jianianhualong* in having a single pneumatic foramen on the lateral aspect of the anterior cervicles, differs from *Sinovenator* in having dorsoventrally compressed sacral centra, differs from *Saurornithoides*, *Troodon*, *Mei*, *Gobivenator*, and *Almas* in having the extremely dorsoventrally compressed sacral centra, differs from all known troodontids in having a ventral groove across the second to the fifth sacral centra (a similar ventral groove is also present in *Zanabazar(Norell et al. 2009)* and *Latenivenatrix*, but it is shorter and inconspicuous in these taxa relative to that present in LH PV39), and differs from *Saurornithoides*, *Zanabazar*, *Latenivenatrix*, and *Gobivenator* among other troodontinaes in having 5 sacral vertebrae. Comparisons with other troodontids, including *Tochisaurus*, *Geminiraptor* (Senter *et al*. 2010), *Philovenator*, *Linhevenator*, and *Borogovia* (Osmólska 1987), are impossible due to the lack or unavailable in these taxa of preserved elements that are equivalent to those known for LH PV39.

The most unusual feature of LH PV39 is the presence of the extremely dorsoventrally compressed sacral centra. Spool-like sacral centra is typical for most theropods including coelophysids (Tykoski 2005), neoceratosaurian (Wang *et al*. 2017), basal tetanuran (Hu 1993), tyrannosaurids (Brochu 2003), ornithomimus (Kobayashi & Lü 2003; Makovicky *et al*. 2004), therizinosaurids (Zanno 2010), most oviraptorosaurs (Balanoff & Norell 2012; Lü & Zhang 2005), troodntids (Shen *et al*. 2017a; Shen *et al*. 2017b) and dromaeosaurids (Norell & Makovicky 1997), whereas dorsoventrally compressed sacral centra have been previously known only in troodontids *Sinovenator* (Xu *et al*. 2002), *Talos* (Zanno *et al*. 2011), *Zanabazar* (Norell *et al*. 2009), and *Latenivenatrix* (van der Reest & Currie 2017); the dromaeosaurids *Rahonavis* (Forster *et al*. 1998), *Buitreraptor* (Novas *et al*. 2018), and *Mahakala* (Turner *et al*. 2007; Turner *et al*. 2011); and are present in the oviraptorosaurs *Microvenator* (Makovicky & Sues 1998) and *Chirostenotes* (Currie & Russell 1988; Sues 1997). The functional implications of this condition remain unclear, though its distribution is limited to Pennaraptora among theropods. Transversely compressed sacral centra are unique to Alvarezsauridae (Chiappe *et al*. 2002), representing an synapomorphy of this clade. Among the taxa whose sacral centra are dorsoventrally compressed, only the centra of *Rahonavis* are as extremely compressed as those seen in LH PV39, but the sacral centra of LH PV39 differs from that of *Rahonavis* in its transversely broad appearance, the presence of a more prominent ventral groove, and absence of pneumatic foramina on the lateral aspects of the centra. In addition, the middle sacrals are wider than the anterior and posterior ones in LH PV39 similar to the condition seen in *Sinovenator*, *Microraptor* and basal birds (Xu *et al*. 2002; Xu *et al*. 2003), whereas in *Rahonavis*, the sacral series becomes transversely narrow posteriorly (Forster *et al*. 1998). Although the sacral centra are spool-like in most theropods, in some troodontids, such as *Zanabazar*, the sacral neural canal can be dorsoventrally compressed when viewed in cross-section but still lesser compressed than that in LH PV39 (Norell *et al*. 2009). It should be noted that the sacral neural canal is also dorsoventrally compressed in LH PV39, and CT images confirm that it is unlikely to be a result of postmortem deformation or erosion. However, the presence and distribution of this condition in other theropods is difficult to determine given that observation of these details is unavailable for most taxa.

*Linheventator* (Xu *et al*. 2011a) and *Philovenator* (Xu *et al*. 2012) are two previously reported troodontids that were collected from the Wulansuhai Formation of Inner Mongolia, but they were not recovered from the same locality of LH PV39. However, little can be said about the vertebral morphology of these taxa because the cervical and sacral vertebrae are not well preserved in *Linhevenator* (Xu *et al*. 2011a), and the entire vertebral column is absent in *Philovenator* (Xu *et al*. 2012). Phylogenetic analyses confirm the troodntinae affinities of all three specimens (Tsuihiji *et al*. 2014; Xu *et al*. 2011a; Xu *et al*. 2012). The Mongolian troodontid *Almas* is recovered as a sister taxon of *Linhevenator* and *Philovenator* in Analyses 2 and 4, suggesting the position of *Almas* seems to be unaffected by character choice, coding discrepancies and taxon sampling (Fig. 5 & 7). The statistically weakly supported phylogenetic analytic results suggest the unstable phylogenetic position of LH PV39 is primarily due to the absence of synapomorpgies, and this is further supported by the unaltered phylogenetic position of *Philovenator* when coded the remains of both LH PV39 and *Philovenator* as a composite OTU (Figs. 5 & 7). The relatively small body size and the complete fused sacral vertebrae suggest LH PV39 is unlikely to represent a juvenile individual of *Linheventator*, but the exact relationship between LH PV39 and *Philovenator* remains unclear due to the lack of overlapping elements from both taxa, pending the recovery of new information. Therefore, current evidence are not strong enough to name LH PV39 as a new taxon or assign it to any known troodontids.

It has been suggested that the evolution of troodontids was dominated by a trend of increased body size, as taxa recovered from the Upper Cretaceous are usually larger than those collected from older sediments (Turner *et al*. 2007; Xu *et al*. 2012). Indeed, troodontids recovered from Lower Cretaceous are chicken sized (Shen *et al*. 2017a; Shen *et al*. 2017b; Xu *et al*. 2017; Xu & Norell 2004; Xu *et al*. 2002), whereas most taxa found from the Upper Cretaceous are usually 1∼2 meters long (Norell *et al*. 2009; Tsuihiji *et al*. 2014; Xu *et al*. 2011a). However, *Philovenator* (Xu *et al*. 2012) and *Almas* (Pei *et al*. 2017b), which was recovered from the Upper Cretaceous of Inner Mongolia, China and the Ukhaa Tolgod, Mongolia, respectively, are two exceptions for their unusual small body sizes with the estimated body length is about 60 cm (Table 2). Another troodontid MPC-D 100/1129 collected from the Upper Cretaceous of Mongolia is also significantly smaller than other Late Cretaceous troodontids as well (Erickson *et al*. 2009), but this specimen has not been systematically described (Norell, personal communication to the senior author), and hence detailed information for comparison remain unavailable.

**Table 2.** Measurements of the maximum femoral length of the selected troodontid specimens.

| Taxon | Specimen number | Maximum<br>Femoral length | Ontogenetic<br>status | References |
| --- | --- | --- | --- | --- |
| *Troodon* | MOR 748 | 483 | adult | (Erickson *et al.* 2009) |
| *Linhevenator* | LH PV21 | 240 | adult | (Xu *et al.* 2011a) |
| *Byronosaurus* | MPC-D 100–984 | 150 | adult | (Erickson *et al.* 2009) |
| *Daliansaurus* | DNHM D2885 | 130.8 | Adult? | (Shen *et al.* 2017a) |
| *Jianianhualong* | DLXH 1218 | 116~117 | adult | (Xu *et al.* 2017) |
| *Liaoningvenator* | DNHM D3012 | 111 | subadult‡ | (Shen *et al.* 2017b) |
| *Philovenator* | IVPP V10597 | 86.5 | subadult | (Xu *et al.* 2012) |
| Unnamed Troodontid | MPC-D 100/1129 | 84 | ? | (Erickson *et al.* 2009) |
| *Mei* | IVPP V12733 | 81 | juvenile | (Xu *et al.* 2002) |
| *Almas* | MPC-D 100–1323 | 80 | subadult | (Erickson *et al.* 2009) |
| *Mei* | DNHM D2154 | 65* | juvenile** | (Gao *et al.* 2012) |

However, we argue that the trend of increased body size during the evolutionary history of troodontids, if present, is not as significant as has previously been suggested. Many Early Cretaceous taxa, such as *Daliansaurus*, *Jianianhualong* and *Liaoningvenator*, are even larger than *Philovenator* and *Almas*, and are presumably larger than LH PV39 (Table 2). Histological analyses suggest the referred specimens of *Mei* (DNHM D3012) and the holotype of *Liaoningvenator* (DNHM D2154) had not reached somatic mature when they perished (Gao *et al*. 2012; Shen *et al*. 2017b), indicating that the mature individuals of these taxa could be even larger. Currently, the effort of reconstructing the growth curve of the known specimens of troodontids seems in vain because many troodontids are represented by fragmentary elements (e.g. *Geminiraptoz* (Senter *et al*. 2010) and LH PV39), and the estimated growth stage of some taxa is unclear (e.g. *Almas*, the histological study of the type specimen has been performed but the growth stage at which it perished has not been published (Erickson *et al*. 2009)). The available histological evidence suggest the small-bodied troodontids may have adopted different growth strategies from those with large body sizes, as the small taxa usually grow slowly and the diaphysis of lone bone is primarily comprised of parallel-fibered bone (e.g. *Philovenator* (Xu *et al*. 2012), note the histological section of *Almas* has been performed though not been published in detailed (Erickson *et al*. 2009)). By contrast, large taxa grow faster and their long bone histological sections are dominated by fibrolamellar bone (e.g. *Troodon* (Varricchio 1993)). This is in consistent with the general growth pattern of maniraptorans as has been previously predicted (Erickson *et al*. 2009), which suggests the size disparity existing within Troodntidae would have been much considerable towards the end of Cretaceous.

## Acknowledgements

We thank team members of the Long Hao Institute of Geology and Paleontology-University of Chicago joint expedition for collecting this specimen, Y. Feng and S. Yuan for CT scanning and 3D segmentation, T. Gao, S. Chen and Q. Lin for helping photography, and X. Xu and C. Forster for allowing the senior author to examine the replicate of *Rahonavis*. S.W. was supported by the National Natural Science Foundation of China (41602013) and HFSP (LT-000728/2018).

## Author contributions

S.W. and L.T. designed research; S.W., Q.T., Q.Z., J.S., H.Z. and L.T. performed the research, S.W. and Q.Z. wrote the paper.

